# Inbreeding depression, heterosis, and outbreeding depression in the cleistogamous perennial *Ruellia humilis*

**DOI:** 10.1101/2023.05.28.542629

**Authors:** Tatyana Y. Soto, Juan Diego Rojas-Gutierrez, Christopher G. Oakley

## Abstract

What maintains mixed mating is an evolutionary enigma. Cleistogamy, the production of both potentially outcrossing chasmogamous, and obligately selfing cleistogamous flowers on the same individual plant, is an excellent system to study the costs of selfing. Inbreeding depression can prevent the evolution of greater selfing within populations, and heterosis in crosses between populations may further tip the balance in favor of outcrossing. Few empirical estimates of inbreeding depression and heterosis in the same system exist for cleistogamous species. We investigate the potential costs of selfing by quantifying inbreeding depression and heterosis in three populations of the cleistogamous perennial *Ruellia humilis* Nutt (Acanthaceae). We performed hand-pollinations to self, and outcross within and between populations, and measured seed number, germination, total flower production, and estimated cumulative fitness for the resulting progeny in a greenhouse experiment.

We found moderate inbreeding depression for cumulative fitness (<30%) in two populations, but outbreeding depression for crosses within a third population (−26%). For between population crosses, there was weak to modest heterosis (11-47%) in two of the population combinations, but modest to strong outbreeding (−21 to −71%) depression in the other four combinations. Neither inbreeding depression nor heterosis was of sufficient magnitude to explain the continued production of CH flowers given the relative energetic advantage of CL flowers previously estimated for these populations. Outbreeding depression either within or between populations makes the maintenance of chasmogamous flowers even harder to explain. More information is needed on the genetic basis of cleistogamy in order to resolve this conundrum.

## INTRODUCTION

The degree of inbreeding influences the amount of genetic diversity within plant populations (Hamrick and Godt, 1996). Inbreeding may thus have consequences for potential adaptation to fluctuating or changing conditions (Glémin and Ronfort, 2013; Hartfield et al., 2017; Hodgins and Yeaman, 2019; Muyle et al., 2020) or population persistence (Coates et al., 2007; Cheptou, 2018). Many angiosperms have flowers that produce both male and female gametophytes. In the absence of a genetic self-incompatibility mechanism, these flowers are capable of self-fertilization, which is the most extreme form of inbreeding (Huenneke, 1991; Jump et al., 2009). Much of the literature on plant mating system evolution has focused on the costs and benefits of selfing. Inbreeding depression is recognized as the primary short-term cost of that can prevent the evolution of greater selfing rates (Lloyd, 1979; Lande and Schemske, 1985; Charlesworth and Charlesworth, 1987; Goodwillie et al., 2005; Charlesworth and Willis, 2009). Inbreeding depression occurs when partially recessive, deleterious alleles are exposed to selection in homozygous progeny produced by selfing, reducing their fitness relative to progeny derived from outcrossed. On the other hand, facultative selfing is expected to have an automatic transmission advantage (Fisher, 1941), a 50% increase in fitness by passing on three copies of gametes (self-fertilization plus donation of pollen to outcrossing) compared to two copies for obligate outcrossers. Much of the early theory on plant mating system evolution predicts that mixed selfing and outcrossing (mixed-mating) is not stable (Lloyd, 1979; Lande and Schemske, 1985; reviewed in Goodwillie et al., 2005; Oakley et al., 2007). Above a 50% threshold of inbreeding depression, the costs of selfing outweigh the benefit and complete outcrossing is favored, below this threshold complete self-fertilization is favored (Lande and Schemske, 1985).

Despite this prediction, there is significant variation in selfing rates within and among species (Goodwillie et al., 2005; Whitehead et al., 2018), and levels of inbreeding depression can vary independently of selfing rate (Winn et al., 2011). This suggests that inbreeding depression and the automatic transmission advantage alone are insufficient to explain the evolution mating systems. For example, mixed-mating is reported for 42% of angiosperms (Vogler and Kalisz, 2001; Goodwillie et al., 2005). In many of these cases of mixed-mating, it is unclear the extent to which selfing is an accidental result of pollen transfer between flowers on the same plant. Such geitonogamy increases with larger floral displays, resulting in a trade-off between increased pollinator attraction and increased selfing rates (Harder and Barrett, 1995; Galloway et al., 2002; Karron et al., 2004).

To evaluate the forces maintaining mixed-mating, it would be ideal to examine systems with specific mechanisms to promote both selfing and outcrossing such as cleistogamy. Cleistogamy is a floral heteromorphism where individual plants produce both potentially outcrossing chasmogamous (CH) flowers and closed obligately selfing cleistogamous (CL) flowers (Kuhn, 1867; Lord, 1981). To clarify, we refer to cleistogamy or cleistogamous to refer to the syndrome of producing both flower types, and cleistogamous (CL) flowers to refer to the selfing flower type (Lord, 1981). Cleistogamy has been widely underutilized in mating system evolution theory (Oakley et al., 2007), likely because of an emphasis on generality (Lande and Schemske, 1985; Goodwillie et al., 2005). While the particulars of this floral heteromorphism do not fit a more general framework, we argue that is a feature and not a bug. Mixed production of CH and CL flowers occurs in over 50 plant families and 220 genera, and has evolved independently multiple times from a CH only ancestral state, with only three documented evolutionary losses of CH flowers (Culley and Klooster, 2007). The repeated independent origins, broad taxonomic occurrence, and persistence of both obligately selfing and potentially outcrossing flowers on the same individual plant provides strong evidence that cleistogamy is an adaptive strategy for mixed-mating (Culley and Klooster, 2007; Oakley et al., 2007).

The production of obligately selfing CL flowers will increase the risk of inbreeding depression in cleistogamous species. However, the selfing rate is expected to co-evolve with inbreeding depression (Lande and Schemske, 1985). This is because recurrent selfing should purge partially recessive, strongly deleterious alleles by exposing homozygous individuals to selection (Uyenoyama and Waller, 1991; Latta and Ritland, 1994). In finite natural populations however, selfing and/or genetic drift can reduce the efficacy of selection against deleterious alleles (Whitlock et al., 2000; Glémin, 2003). Because new mutations are on average mildly deleterious and partly recessive (Eyre-Walker and Keightley, 2007) an accumulation of deleterious alleles fixed by drift over time is expected for alleles that are not purged by selection or drift (Whitlock et al., 2000; Glémin, 2003; Charlesworth, 2018). These fixed deleterious alleles cannot contribute to inbreeding depression within populations because they do not segregate between progeny derived from selfing versus outcrossing. However, they will still have negative fitness consequences for all individuals in the population, and therefore present another potential cost of selfing (Byers and Waller, 1999).

To determine if potentially low empirical estimates of inbreeding depression are due to an absence vs. fixation of deleterious recessive alleles, we need to estimate both inbreeding depression and heterosis. Heterosis is important in crops like maize, where F_1_ hybrids have greater yield than inbred parental lines (Lippman and Zamir, 2007). In natural populations, greater fitness of crosses between-comparted to within-populations can be used to infer dominance complementation of fixed deleterious recessive alleles (Crow, 1948; Whitlock et al., 2000). Heterosis is indeed common in wild populations (Sheridan and Karowe, 2000; Marr et al., 2002; Weller et al., 2005; Busch, 2006; Anderson and Hedgecock, 2010). Empirical evidence of greater heterosis in small or highly selfing populations (Paland and Schmid, 2003; Oakley and Winn, 2012; Oakley et al., 2015a; Charlesworth, 2018) is consistent with the fixation of partially recessive deleterious alleles contributing the the genetic basis of heterosis.

Heterosis has also been proposed as a potential mechanism to maintain CH flowers in cleistogamous species (Oakley et al., 2007). Here, selfing via CL flowers provide a greater energetic economy, and progeny derived from CL selfing often have comparable fitness to progeny derived from CH outcrossing. Hence, additional forces seem necessary to explain the evolutionary maintenance of CH flowers, and heterosis in rare outcrossing between populations is one potential explanation for such an advantage (Oakley et al., 2007). On the other hand, outbreeding depression, the reduction in fitness of progeny from crosses between populations relative to crosses within populations, is also possible (Lynch, 1991; Schierup and Christiansen, 1996; Edmands, 1999). Such outbreeding depression is increasingly found to be common in between population crosses in highly selfing species (Volis et al., 2011; Gimond et al., 2013; Oakley et al., 2015a; Oakley et al., 2019; Clo et al., 2021), and would likely select for increased selfing.

Quantifying both inbreeding depression and heterosis is an important step for understanding the effects of mating system on patterns of deleterious recessive variation. It is additionally important in understanding the maintenance of CH flowers in cleistogamous species. Unfortunately, estimates of inbreeding depression and heterosis in the same system are uncommon (but see for e.g, Paland and Schmid, 2003; Weller et al., 2005; Oakley and Winn, 2012; Oakley et al., 2019), and to our knowledge such estimates are only available for one cleistogamous species (Seguí et al., 2021). Additionally, few studies in cleistogamous species estimate inbreeding depression independent of differences between flower type (reviewed in Oakley et al., 2007), but see (Culley, 2000; Munguía-Rosas et al., 2013; Ansaldi et al., 2019). Here we used a greenhouse study to estimate multiple fitness components and cumulative fitness for progeny from controlled pollinations to calculate both inbreeding depression and heterosis (or outbreeding depression) in three populations of the cleistogamous perennial *Ruellia humilis* Nutt (Acanthaceae). Our primary questions are: 1) Is inbreeding depression within populations greater than the 50% threshold required to maintain CH outcrossing? 2) Do crosses between populations result in heterosis or outbreeding depression? 3) If crosses between populations result in heterosis, is the magnitude sufficient to provide enough of an advantage to explain their evolutionary maintenance?

## MATERIALS AND METHODS

### Study system

*Ruellia humilis* Nutt (Acanthaceae) is a short-lived herbaceous perennial native to prairies and sandy soils throughout the midwestern United States (Fernald, 1945). It is part of the humilis clade that includes the five spp. native to eastern North America (Tripp, 2007). All species in this clade are cleistogamous, with individual plants producing both chasmogamous (CH) flowers and cleistogamous (CL) flowers. In *R. humilis,* selfing occurs via CL flowers, and CH flowers are self-compatible and occasionally self-fertilize. The proposed mechanism for autogamous selfing in CH flowers is that the anthers (stamens fused to the corolla tube) brush against the stigma as the corolla tube abscises (Long and Uttal, 1962). It has been suggested that both flower types are produced simultaneously over the entire flowering period from June to September (Long and Uttal, 1962). Hawkmoths are the primary pollinators of *R. humilis* with occasional visits from long-tongued bees and butterflies (Heywood et al., 2017). Anther dehiscence of both CH and CL flowers occurs after dusk and CH flower anthesis occurs a couple of hours later. Both flower types last for less than one day. Seed dispersal in this species is achieved by explosively dehiscing fruits, dispersing seeds as far as 7m away from the maternal plant (Heywood et al., 2017; Cooper et al., 2018).

Because of widespread loss of prairie habitat in the midwestern United States, a limited number of remnant native populations remain. We sampled three natural populations in Indiana and Illinois (**Appendix S1**). Sand Ridge (SR) and St. Mary’s (SM) are both populations found in cemetery prairie remnants in Central Indiana. Wilmington (WN) is a roadside population located in Northeastern Illinois and is located furthest from the other populations. All three populations are subjected to frequent mowing during the flowering period. Sand Ridge was the largest of the three populations and we noted abundant seed predation at the time of collection. St. Mary’s is the smallest of the three populations and is likely subjected to the most frequent mowing.

### Maternal line seed collection and germination

We collected seed from an average of 23 maternal lines (range: 19-25) from each of the three populations in the Fall of 2020. Seed was cold stratified on damp paper towels in the dark in a refrigerator (∼4°C) for 8-10 weeks to break dormancy (Baskin and Baskin, 1998). After cold stratification, one seed per cell was sown in 200 cell flat inserts (cell volume = 14 cm^3^, Greenhouse Megastore Danville, IL) filled with a 2:1 mixture of MetroMix potting soil (Sun Gro Horticulture Agawam, MA) and sand. Flats were placed in an incubator (Percival Scientific, Perry, IA) set at 28°C with 13-hour days at the maximum photosynthetic active radiation setting (65 μmoles/m^2^/sec) until the seeds germinated. A total of 256 of the resultant seedlings were transplanted into 9cm pots (volume = 245 cm^3^, Greenhouse Megastore Danville, IL) filled with the same media mixture two weeks after sowing. Five weeks later, seedlings were again transplanted into 15cm (volume = 1556 cm^3^, Greenhouse Megastore Danville, IL) pots with a 1:3 mixture of topsoil and sand. This mixture was selected to be well-draining and similar to soils supporting *R. humilis* populations. Plants were watered as needed and grown with 12-hours of supplemental lighting per day in a heated greenhouse (minimum: 20°C).

### Hand pollination design and seed collection

To quantify inbreeding depression and heterosis, we performed four types of hand pollinations on CH flowers for each of the three focal populations for a total of twelve progeny types (**Appendix S2**). Hand pollinations were performed on plants from 21, 18, and 23 maternal lines from SR, SM, and WN respectively. These cross types included self-fertilization (SELF), outcrosses within populations (WIN), and crosses between population (BET) for each pairwise combination of populations. Hand pollinations were initiated within 2 weeks of when plants began to flower in April 2021 and were completed by July 2021. Each evening, pollen and ovule donors were selected based on which plants would flower the next day. For all pollinations, we carefully removed the immature anthers from the ovule donor using forceps and scissors. The following morning, dehisced anthers were removed from the pollen donors, and we applied the pollen to the stigmas of the ovule donors. Forceps and scissors were sterilized using 70% ethanol and rinsed with DI water between each emasculation and pollination. We confirmed that the emasculation procedure did not result in accidental self-pollination by performing 60 control emasculations (emasculated but not pollinated), none of which produced seed. In total, we performed 1,981 hand pollinations, 1,640 of which successfully set fruit. Fruits were collected in autumn 2021 once the fruits had turned brown and were close to dehiscing. Fruits were placed individually in coin envelopes and allowed to dehisce prior to counting seed number per fruit.

### Fitness assay

Once all fruit was matured and collected, we counted seed number per fruit and scored the probability of germination of these seeds. We quantified seed number per fruit for a subset of 667 fruits from successful crosses (**Appendix S3**). Seeds were then cold stratified following the protocols described above, pooling seeds from fruits that shared the same unique combination of maternal and paternal lines. After stratification we selected between 110-118 seeds for each of the 12 progeny types (1,710 seeds in total) to assay germination (**Appendix S3**). These seeds were sown into 288-cell flat inserts (volume = 7 cm^3^, Greenhouse Megastore, Danville, IL) filled with a 2:1 mixture of Berger BM2 potting soil (Berger Horticulture Products, Sulphur Springs, TX) and sand. Seeds were sown individually in cells in a randomized block design with a total of 6 blocks (one block = one 288-cell insert), each consisting of 19 seeds from each of the 12 cross types. Remaining seeds were sown in a fully randomized design into 4 additional inserts using the same soil and sand mixture to supplement later life stages of the fitness assay in case of low rates of germination. All 10 flats were placed in an incubator set at conditions described previously, until the seeds germinated. Each germinated seed was scored as 1, and ungerminated seeds were scored as 0.

After germination, plants were grown in a greenhouse to measure probability of reproduction and total number of flowers per reproductive plant. Three weeks after sowing, 1,216 seedlings with two sets of true leaves were transplanted into 9 cm pots filled with a 2:1 ratio of Berger BM2 potting soil and sand. Each seedling was planted into a completely randomized array. Due to greenhouse space constraints, a subset of 773 seedlings were retained for the remainder of the experiment. Once these seedlings had four sets of true leaves, they were transplanted in a fully randomized fashion into their final pot size (10.79cm x 10.79cm x 12.38cm, Greenhouse Megastore, Danville, IL) filled with the same sand/soil mixture and fertilized with Osmocote Plus (Bloomington, IN) following the manufacturer recommended application. Due to differential germination among progeny types, the final design to measure subsequent fitness components was unbalanced (**Appendix S3**). Because of low mortality, we scored a combined probability of survival and reproduction using 0 and 1, for plants that did not flower or flowered respectively. Plants began flowering in April 2022 and we estimated the total number of flowers produced per individual in December 2022 by counting the persistent calyces.

### Statistical analysis and calculation of inbreeding depression and heterosis

For each fitness component, we tested for the effect of progeny type as a fixed effect using ANOVA. Progeny type is a composite effect that includes both maternal population (3 levels) and pollination type (4 levels: SELF, WIN, and 2 levels of BET with different paternal populations) for a total of 12 levels. For seed number, we used a normal error distribution based on visual inspection of the residuals. For probabilities of germination and reproduction, we used generalized linear models with a binomial error distributions and logit link functions. For probability of germination (the only stage with blocking), we also included an effect of block as a fixed effect to account for spatial variability. Block was treated as a fixed effect because the small number of blocks preclude them from being truly random sample (Bennington and Thayne, 1994). For flower number, we used a generalized linear model with a Poisson error distribution and a log link function. For each analysis, if the overall effect of progeny type was significant, we performed pairwise contrasts for three *a priori* hypotheses for each population. To investigate inbreeding/outbreeding depression within populations, we contrasted selfed vs. within population crosses. To investigate heterosis/outbreeding depression between populations, we contrasted within population cross vs. between population crosses separately for both possible pollen donors.

For individual fitness components we calculated population level inbreeding depression as relative performance (Ågren and Schemske, 1993) because of populations with negative inbreeding depression. We express this measure as a percentage, and it is bounded by −100% and 100% and quantifies the proportional difference between WIN and SELF [(WIN-SELF)/max(WIN, SELF)]*100 relative to the most fit progeny type. Heterosis and outbreeding depression were calculated as the percent increase/decrease in fitness of between population crosses relative to within population crosses [(BET– WIN)/WIN]*100 separately for each of the two BET categories for the two potential paternal populations.

To estimate cumulative fitness (total flower production in the next generation from an initial pollination event) of each of the twelve progeny types, we multiplied population level means for all 4 fitness components. We then calculated cumulative estimates of population level inbreeding depression, heterosis, and outbreeding depression as above, and tested if these estimates were significantly different from zero using 95% confidence intervals (CI) derived from bootstrap resampling. For each of the 12 progeny types, we resampled in order to obtain 60 measures of cumulative fitness, roughly corresponding to the number of fruits per progeny type used to initiate the experiment. For each measure (60 measures x 12 progeny types), we randomly drew one value from each of empirically determined fitness components for a given population and pollination type and multiplied them together, repeating this procedure 125 times to get a mean bootstrapped estimate of cumulative fitness. The 125 replicates for the bootstrapping is arbitrary, but was chosen to be similar to the largest sample size of adult plants for any progeny type included in the fitness assay. This stratified sampling was necessary because of the large number of cumulative fitness estimates of zero for one or more progeny types, driven by low probability of germination, particularly in SM. These zero estimates are realistic, but make calculating ratios impossible (i.e. dividing by zero is undefined). These 60 mean bootstrapped estimates of cumulative fitness for each progeny type were then used to calculate means and standard errors inbreeding depression, heterosis, and outbreeding depression as above. We calculated a 95% CI based on these means and standard errors (assuming a normal distribution) around population level estimates of inbreeding depression, and heterosis or outbreeding depression.

## RESULTS

Overall, seed number per fruit was similar across populations (**Fig. 1, Appendix S3**), as was probability of reproduction. There was more variation among populations in overall probability of germination, which was high in WN, very low in SM, and intermediate in SR (**Fig. 1, Appendix S3**). Despite the low germination in SR, this population had the greatest overall flower production, with SM and WN both producing fewer flowers on average (**Fig. 1, Appendix S3**). Our primary interest was the effect of progeny type, and this had a significant effect on all fitness components except probability of reproduction (**Table 1**).

**Figure 1.**
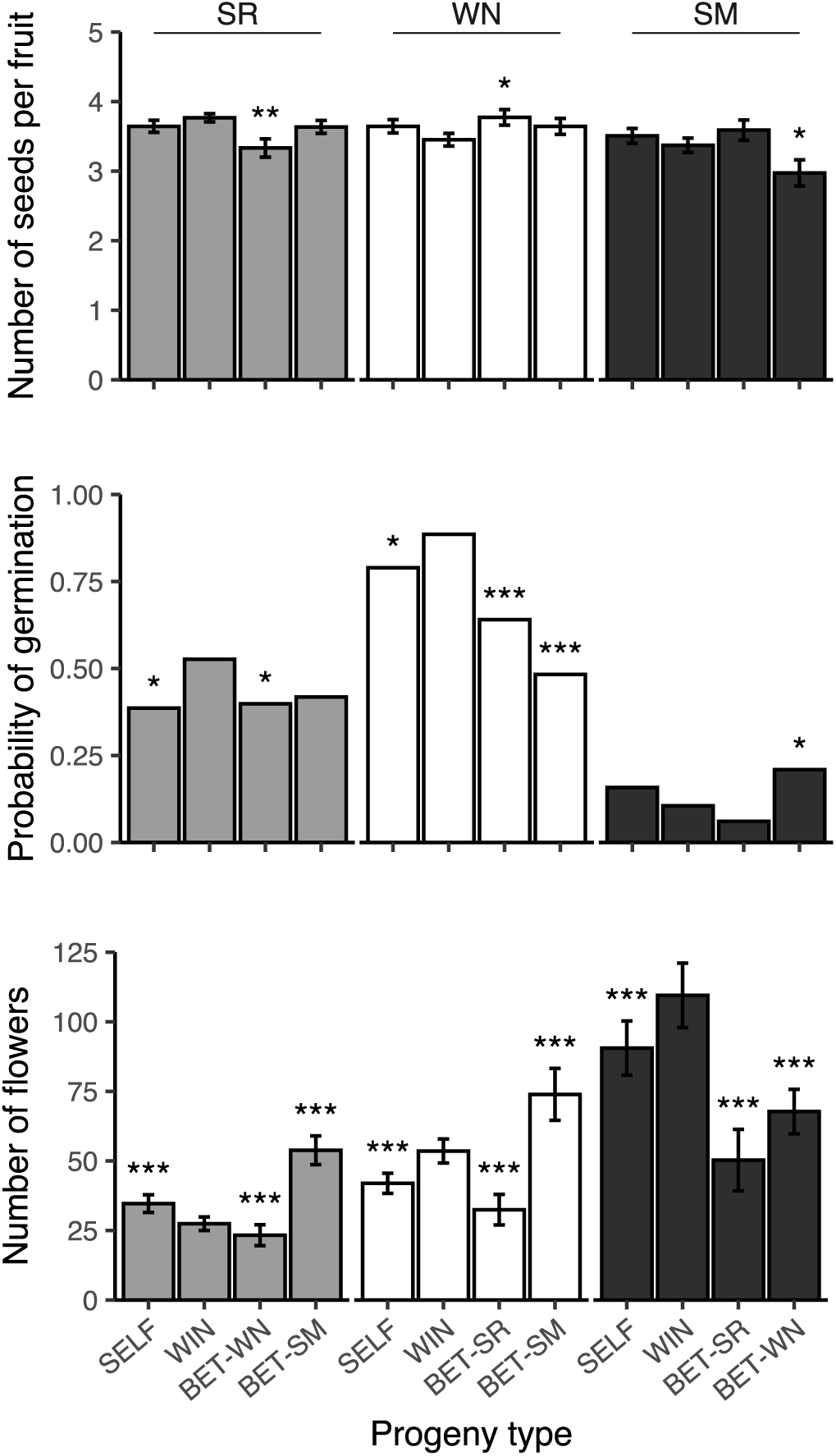
Mean fitness components for the twelve progeny types (four cross types for three populations). Error bars are ± SE. Population after the dash for the BET crosses indicates the pollen donor population. Asterisks indicate significant contrasts for *a priori* hypothesis tests (* p < 0.05, ** p < 0.01) for each cross type in each population to the WIN cross type (See Table 2 for details).

**Table 1.** ANOVA results for the effect of progeny type on fitness components. F-ratios, degrees of freedom (numerator, denominator), and p-values are given for seed number per fruit as this was the only fitness component analyzed with a normal error distribution. For the remaining fitness components analyzed with binomial (probabilities of germination and reproduction) and Poisson (flower number) error distributions, Chi square values, df, and P values are given. Germination was the only stage with a blocked design.

| **Fitness component** | **Progeny type** |  |  | **Block** |  |  |
| --- | --- | --- | --- | --- | --- | --- |
| | F or $\chi^2$ | df | P | $\chi^2$ | df | P |
| Seed number per fruit | 3.8 | 11, 656 | <0.001 |  |  |  |
| Probability of germination | 477 | 11 | <0.001 | 29 | 5 | <0.001 |
| Probability of reproduction | 19 | 11 | 0.059 |  |  |  |
| Flower number | 7015 | 11 | <0.001 |  |  |  |

**Table 2.** Results of linear contrasts from the above ANOVA models on fitness components testing three *a priori* hypotheses for each of the three populations. Comparisons are between self-pollinations (SELF), outcrosses within populations (WIN), and crosses between populations (BET-) where the population after the dash indicates the pollen donor population. Test statistics are t for fitness components with normal error distributions or Chi square otherwise. No contrasts were performed for probability of reproduction because there was no significant effect of progeny type in the overall ANOVA (see table 1).

| Pop. | Comparison | Seed number |  | Probability of |  | Number of |  |
| --- | --- | --- | --- | --- | --- | --- | --- |
|  |  | per fruit |  | germination |  | flowers |  |
| | | t | P | $\chi^2$ | P | $\chi^2$ | P |
| SR | SELF vs. WIN | -0.9 | 0.324 | 4.6 | 0.031 | 95 | <0.001 |
|  | BET-WN vs. WIN | 2.9 | 0.004 | 3.8 | 0.048 | 24 | <0.001 |
|  | BET-SM vs. WIN | 0.9 | 0.369 | 2.7 | 0.097 | 761 | <0.001 |
| WN | SELF vs. WIN | 1.4 | 0.171 | 3.9 | 0.044 | 138 | <0.001 |
|  | BET-SR vs. WIN | 2.1 | 0.038 | 20 | <0.001 | 320 | <0.001 |
|  | BET-SM vs. WIN | -1.3 | 0.212 | 47 | <0.001 | 215 | <0.001 |
| SM | SELF vs. WIN | 0.9 | 0.329 | 1.4 | 0.235 | 69 | <0.001 |
|  | BET-SR vs. WIN | 1.4 | 0.172 | 1.6 | 0.211 | 359 | <0.001 |
|  | BET-WN vs. WIN | 2.5 | 0.012 | 4.7 | 0.029 | 271 | <0.001 |

### SELF vs. WIN

Significant contrasts between SELF and WIN cross types for a given population are consistent with inbreeding (or outbreeding) depression within a given population. For seed number per fruit, there were no significant contrasts for any of the three populations and estimates of relative performance were small, ranging from −5% to 3% (**Fig. 1, Tables 2 & 3**). For probability of germination, contrasts between SELF and WIN were significant for the SR and WN populations, with estimates of relative performance of 26% and 11% respectively (**Fig. 1, Tables 2 & 3).** Relative performance for germination in SM was moderately large and negative (−39%), but the contrast was not significant for this population, possibly because of the very low overall probability of germination. No contrasts were performed for probability of reproduction because of a lack of a significant overall effect of progeny type (**Table 1**). For number of flowers per reproductive plant, contrasts between SELF and WIN were highly significant for all populations (**Fig. 1, Tables 2 & 3).** Relative performance was positive (i.e. inbreeding depression) for both WN (21%) and SM (17%), but negative (i.e. outbreeding depression for crosses within the population) for SR (−20%). For cumulative fitness (**Fig. 2, Table 3**) we observed significant but modest inbreeding depression for SR (14%) and WN (27%). There was significant outbreeding depression for within population crosses for SM (−26%). Across the life cycle, we found a mixture of inbreeding and outbreeding depression for different fitness components and cumulative fitness (**Table 3**). The direction of the effect was most consistent for WN where inbreeding depression was observed for germination probability, flower number, and cumulative fitness.

**Figure 2.**
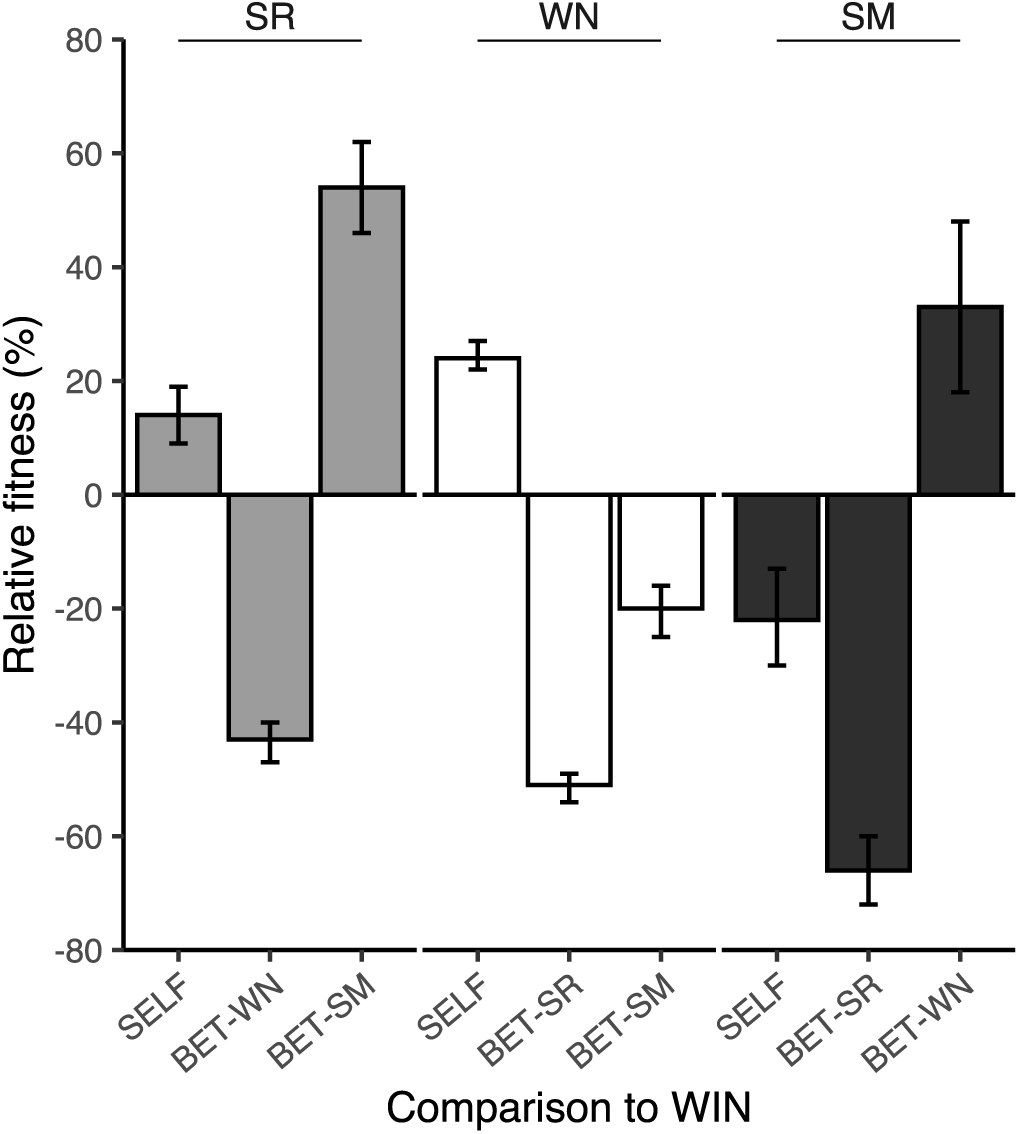
Bootstrapped estimates of mean and 95% confidence intervals for inbreeding/outbreeding depression and heterosis. Bars represent comparisons between WIN and each of the three other cross types for each population. SELF vs. WIN indicates inbreeding (positive values) or outbreeding (negative values) depression. The two comparisons of BET to WIN (with both possible pollen donor populations) indicate heterosis (positive values) or outbreeding depression (negative values).

**Table 3.** Population level estimates of percent inbreeding/outbreeding depression and heterosis for cumulative fitness and fitness components for the three populations. In all cases, negative values indicate outbreeding depression for crosses either within (comparisons with SELFs), or between (comparisons with BET) populations. Positive values for comparisons with SELF and BET indicate inbreeding depression and heterosis respectively.

| **Pop.** | **Comparison** | **Cumul.**<br>**fitness** | **Seed num.**<br>**per fruit** | **Germ.**<br>**prob.** | **Repro.**<br>**Prob.** | **Flower**<br>**num.** |
| --- | --- | --- | --- | --- | --- | --- |
| **SR** | SELF vs. WIN | 14 | 3 | 26 | 1 | -20 |
|  | BET-WN vs. WIN | -50 | -11 | -27 | -12 | -15 |
|  | BET-SM vs. WIN | 47 | -3 | -17 | 1 | 96 |
| **WN** | SELF vs. WIN | 27 | -5 | 11 | 1 | 21 |
|  | BET-SR vs. WIN | -54 | 9 | -28 | -5 | -39 |
|  | BET-SM vs. WIN | -21 | 5 | -45 | 1 | 38 |
| **SM** | SELF vs. WIN | -26 | -3 | -39 | 0 | 17 |
|  | BET-SR vs. WIN | -71 | 6 | -19 | 4 | -54 |
|  | BET-WN vs. WIN | 11 | -12 | 126 | 4 | -38 |

### BET vs. WIN

Significant contrasts between WIN and BET cross types for a given pair of populations are consistent with heterosis (positive values) or outbreeding depression (negative values). For mean seed number per fruit, 3 of the 6 contrasts of within compared to between population crosses were significant (**Fig. 1, Table 2**). The cross to WN using pollen from SR resulted in heterosis (9%), and both crosses with WN as the pollen donor resulted in outbreeding depression, −11% in SR and −12% in SM (**Table 3**). For probability of germination 4 out of the 6 contrasts of WIN and BET were significant, with a suggestive result for a 5^th^ contrast (**Fig. 1, Table 2).** Of these contrasts, there was only a single case of heterosis (126%), in SM with WN as the pollen donor. All other contrasts indicated outbreeding depression ranging from −17% to −45% (**Table 3**). No contrasts were performed for probability of reproduction because of a lack of a significant overall effect of progeny type (**Table 1**). For number of flowers, all six contrasts between WIN and BET were significant (**Fig. 1, Table 2**). There were two cases of heterosis for this fitness component, both involving SM as the pollen donor, 96% in SR and 38% in WN (**Table 3**). The other four cases of outbreeding depression ranged from −15% to −54% (**Table 3**).

For cumulative fitness, all estimates of heterosis and outbreeding depression were significant (**Fig. 2**). There were two cases of heterosis, in SR with pollen from SM (47%) and in SM with pollen from WN (11%). The remaining cases were outbreeding depression (**Table 3**). Both crosses with WN as the ovule donor resulted in outbreeding depression of −54% and −21% with pollen from SR and SM respectively. The other two cases of outbreeding depression for cumulative fitness were SR x WN (−50%) and SM x SR (−71%). Across the life cycle, there was consistent outbreeding depression for all fitness components and cumulative fitness for the SR x WN cross (**Table 3).** For the other five combinations we found a mixture of effects. Three of these (heterosis for one, outbreeding depression for the other two), exhibited consistent effects across germination probability, flower number, and cumulative fitness (**Table 3**). Discordant effects across the life cycle were observed for SM x WN which had 126% heterosis for germination probability, but −38% outbreeding depression for flower number, with a net effect of 11% heterosis overall. To a lesser degree, such differences were also observed for WN x SM which had −45% outbreeding depression for germination probability, but 38% heterosis for flower number, and −21% outbreeding depression overall.

## DISCUSSION

Inbreeding depression is thought to be a central mechanism in preventing the evolution of greater selfing within populations, and heterosis in crosses between populations may further tip the balance in favor of outcrossing. However, there are few empirical estimates of inbreeding depression and heterosis in the same system, particularly for cleistogamous species. We found that levels of inbreeding depression, heterosis, and outbreeding depression in three populations of *Ruellia humilis* vary depending on life stage and ovule/pollen donor population. Cumulative inbreeding/outbreeding depression was significant within all populations and ranged from −26% to 27%. Lastly, we documented two cases of significant heterosis (11% and 47%) and four cases of significant outbreeding depression (−21% to −71%).

### SELF vs. WIN

Our findings indicate modest inbreeding depression in two populations (14 % in SR and 27% in WN) and outbreeding depression within another population (−26% in SM). Weak inbreeding depression fits the expectation for small and/or highly selfing populations, where strongly deleterious alleles may have been purged from the population via some combination of selection or drift (Lande and Schemske, 1985; Glémin, 2003; but see Winn et al., 2011). Outbreeding depression within populations has also been reported in the cleistogamous *Viola jaubertiana* (Seguí et al., 2021), and other highly selfing species (Paland and Schmid, 2003; Volis et al., 2011; Oakley and Winn, 2012; Gimond et al., 2013; Clo et al., 2021). Such outbreeding depression within population outcrossing suggests fixation of alleles within inbred genetic lineages that result in genetic incompatibilities when introduced into the genetic background of other linages via outcrossing within the population. It is unclear why cumulative outbreeding depression occurs only in SM, but this is the smallest of the other three populations. In all cases, inbreeding depression is not sufficiently strong (i.e., not greater than 50%) to explain the evolutionary maintenance of CH outcrossing. This is particularly true because of the greater energetic economy of selfing via CL flowers in this species (Soto et al., 2023), a pattern commonly found in many other cleistogamous species as well (Oakley et al., 2007).

An additional expectation for selfing species is that inbreeding depression should be weak in early life history stages due to purging, but increase across the life cycle because of the cumulative effects of many mildly deleterious alleles (Husband and Schemske, 1996; but see Winn et al., 2011). To some extent, WN fits this pattern with weak inbreeding depression for germination and flower number resulting in somewhat greater inbreeding depression for cumulative fitness. Patterns across life history stages were idiosyncratic for the other two populations. We observed inbreeding depression for germination and outbreeding depression for flower number in SR but found the opposite pattern for SM. Outbreeding depression in early life stages and heterosis in later life stages was also documented in *Medicago trunculata* (Clo et al., 2021).

An important caveat to our estimates of inbreeding depression is that greenhouse studies may underestimate inbreeding depression by eliminating abiotic and biotic stresses typical of the natural environment (Dudash, 1990; Cheptou and Donohue, 2011). However, many examples of environmental dependence of inbreeding depression are cases where the magnitude, but not the sign, of inbreeding depression changes across environments (Oakley and Winn, 2008; Munguía-Rosas et al., 2013; Stojanova et al., 2020). One study of two cleistogamous species reported negative inbreeding depression in a more ‘benign’ environment, but inbreeding depression did not exceed 50% in a more ‘stressful’ environment (Eckstein and Otte, 2005). Additionally, our estimates of inbreeding depression are consistent with estimates in other cleistogamous species (Culley, 2000; Eckstein and Otte, 2005; Oakley and Winn, 2008; Winn and Moriuchi, 2009; Munguía-Rosas et al., 2013; Ansaldi et al., 2019), which are all lower than 50%. Thus, it seems unlikely that field estimates of inbreeding depression in natural populations of *R. humilis* would dramatically change our qualitative conclusions.

### BET vs. WIN

Heterosis for cumulative fitness was found in two of the six between population crosses (11% for SM x WN and 47% for SR x SM). This increased fitness of crosses between relative to crosses within population suggests dominance complementation of different deleterious recessive alleles that have been fixed within populations by drift (Whitlock et al., 2000; Paland and Schmid, 2003; Oakley and Winn, 2012; Lohr and Haag, 2015; Oakley et al., 2015b; Charlesworth, 2018). Moderate heterosis for cumulative fitness has also been reported in the only existing published estimate of heterosis in a natural population of a cleistogamous species (Seguí et al., 2021). As with inbreeding depression, estimates of heterosis across life history stages were idiosyncratic for populations with heterosis for cumulative fitness. For SR x SM, weak outbreeding depression was observed for germination probability, but there was strong heterosis for number of flowers. For SM x WN, the opposite pattern was observed, with strong heterosis for germination and modest outbreeding depression for flower number. Variation in heterosis and outbreeding depression across different life stages has been reported in *Physa acuta* (Escobar et al., 2008).

Strong heterosis for specific fitness components challenges the notion that heterosis is caused by many mildly deleterious alleles, where cumulative heterosis would be expected to be the cumulative effect of weak heterosis at multiple fitness components. Given the idiosyncrasies in relative fitness across life stages, it could be that heterosis at individual stages may be a balance between positive effects of heterosis and negative effects of outbreeding depression. An alternative explanation is that heterosis in this system is caused by more strongly deleterious alleles. Regardless, even the greatest overall estimates of heterosis of 47% are likely insufficient to explain the maintenance of CH flowers both because of the energetic economy of CL selfing, as well as a limited potential for frequent gene flow between populations (Soto et al., 2023).

Outbreeding depression for cumulative fitness was found in four of the six between population crosses. This outbreeding depression was modest (−21%) for one cross, but strong (between −50% and −71%) for the other three crosses. There are no reported estimates of outbreeding depression in natural populations of a cleistogamous species, but to our knowledge only one published study has the relevant data (Seguí et al., 2021). Outbreeding depression is most commonly observed in the F_2_ and later generations, where negative epistatic interactions are thought to play a primary role (Lynch, 1991; Schierup and Christiansen, 1996; Edmands, 1999; Fenster and Galloway, 2000). There is a growing body of literature indicating that outbreeding depression in the F_1_ generation is common for highly selfing species (Volis et al., 2011; Gimond et al., 2013; Oakley et al., 2015a; Oakley et al., 2019; Clo et al., 2021).

Outbreeding depression in *R. humilis* may be exacerbated by habitat loss and fragmentation due to the widespread conversion of prairies to agriculture. These habitat modifications could both increase the chance of fixation of alleles contributing to outbreeding depression in small remnant populations (Barmentlo et al., 2018), and may reduce pollinator movement among populations (Rathcke and Jules, 1993). Indeed, reduced pollinator visitation in *Ruellia humilis* by the primary pollinators (hawkmoths) has been documented (Heywood et al., 2022). Regardless, outbreeding depression in between population crosses cannot be a force that maintains CH flowers in this system.

## CONCLUSIONS

Within populations, we found significant inbreeding depression in two populations, but the magnitude was considerably less than the 50% threshold required to prevent the evolution of further selfing. Outbreeding depression in within population outcrosses for the third population would be expected to exert selection for increased selfing. Outcrossing could also provide a fitness advantage to CH flowers if there is heterosis in rare outcrossing between populations, however we found weak to modest heterosis, and only in a few of the possible population combinations. For the majority of combinations, we found modest to strong outbreeding depression. Taken together we find a fitness disadvantage of CH outcrossing in most cases. When there was an advantage of CH outcrossing, the effect was not strong. Based on a recent study (Soto et al., 2023) quantifying the relative energetic economy of CL selfing in these three populations, there needs to be a 4-10 fold short-term fitness advantage of CH flowers to explain their continued production. Having exhausted the possible proposed explanations for the maintenance of CH flowers, it time to consider other alternative explanations. For example, it could be the loss of CH flowers is genetically constrained by pleiotropic effects of genes on both flower types. It may be feasible to test this hypothesis using artificial selection in species with a rapid life cycle. Additional work on the genetic mechanisms underlying the developmental switch between the two flower types may additionally shed more light on this question.

## Acknowledgments

The authors thank Bob Easter from NICHES Land Trust for allowing us to collect seed from Sand Ridge. We also would like to thank Gordon McNickle for helpful comments on an earlier version of this manuscript, and Nicholas Ryan for assistance with the greenhouse experiment. This work was funded in part by USDA Hatch grant 7000281 to C.G.O via Purdue College of Agriculture, and a Purdue ARGE assistantship to T.Y.S.

## Author Contributions

T.Y.S and C.G.O conceived of and designed the study; T.Y.S conducted the experiment and collected the data; T.Y.S and J.D.R.G. analyzed the data; J.D.R.G. and T.Y.S. produced the tables and figures; T.Y.S drafted the manuscript with help from C.G.O. All authors contributed to revising the manuscript.

## Data Availability Statement

Upon acceptance all raw data will be archived at PURR, the Purdue public data archiving repository.

---

**Appendix S1.** Approximate census sizes of the three focal populations, along with the geographic coordinates and geographic distances between populations (km).

| **Population** | **Census size** | **Coordinates** | **Distance to**<br>**Sand Ridge**<br>**(km)** | **Distance to**<br>**Wilmington**<br>**(km)** |
| --- | --- | --- | --- | --- |
| **Sand Ridge** | 400 | 40.4085° N,<br>86.9665° W | - | - |
| **Wilmington** | 250 | 41.3124° N,<br>88.1321° W | 143 | - |
| **St. Mary's** | 50 | 40.3952° N,<br>86.9172° W | 5 | 168 |

**Appendix S2.**
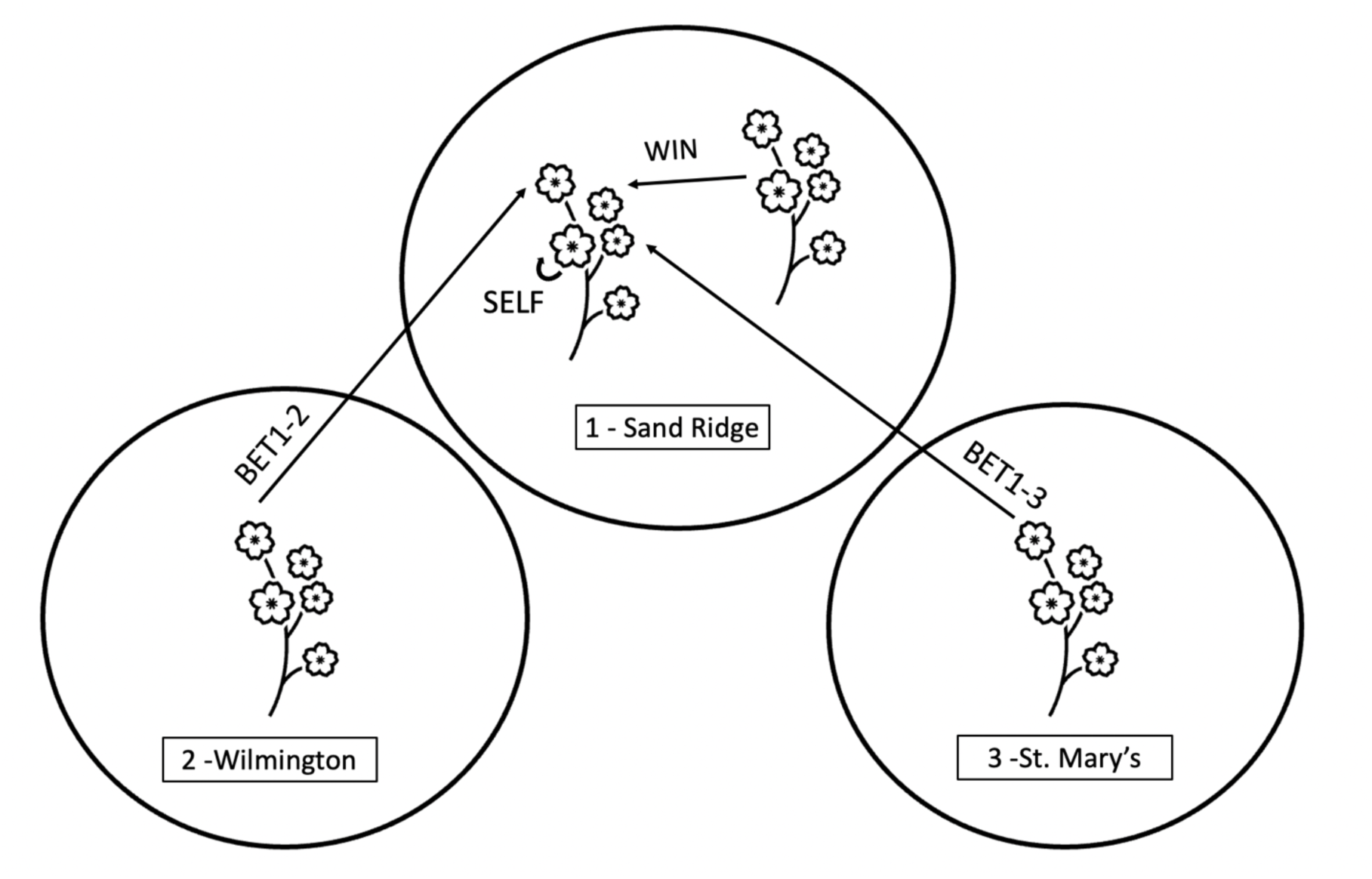
Example pollination design for one plant in one focal population. This design was repeated multiple times for 18-23 maternal lines per population. SELF indicates CH flowers that were self-pollinated. WIN indicates CH flowers that were pollinated with pollen from another maternal line from the same population. BET indicates a CH flower that received pollen from either one of the other two populations, which are associated with a number i.e., BET1-2 is Sand Ridge x Wilmington. Inbreeding depression was calculated by comparing fitness estimates of progeny from SELF and WIN crosses. Heterosis and/or outbreeding depression was calculated by comparing fitness estimates of progeny from WIN and BET separately for each unique combination of maternal and paternal populations.

**Appendix S3.**
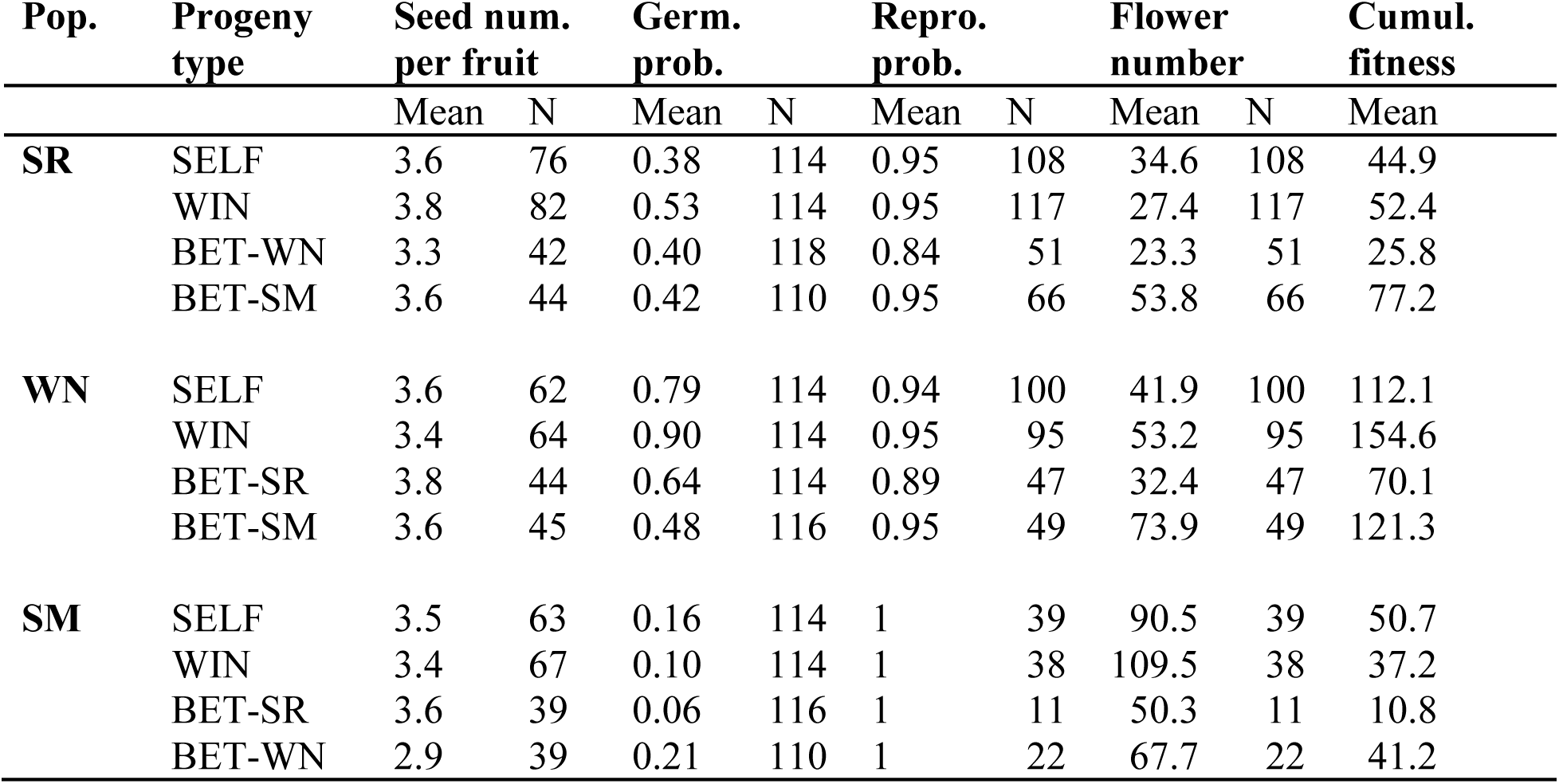
Means and sample sizes for each fitness component and mean cumulative fitness for each of the 12 progeny types (SELF = selfed, WIN = outcrossed within populations, and two BET = outcrossed between populations, with population after the dash indicating the pollen donor population. Sample sizes are the number of fruits counted, the number of seeds sown, the number of juvenile plants censused, and the number of reproductive plants censused, respectively for seed number per fruit, probability of germination, probability of reproduction, and number of flowers respectively.

